# Inotropic effects of 3-hydroxybutyrate in a rodent heart: pressure-volume and isolated rat ventricular trabeculae analysis

**DOI:** 10.64898/2026.09.15.751359

**Authors:** Jan Kropáček, Thu Thao Nguyen, Ema Hájková, Matej Molnár, Luca Monzo, Matúš Miklovič, Vojtěch Melenovský

## Abstract

An administration of 3 hydroxy-butyrate (3-OHB) in experimental animals or humans is associated with an increase of cardiac output (CO), but whether this hemodynamic benefit reflects direct myocardial inotropy or heart unloading due to peripheral vasodilation remains unclear. It is also unknown if myocardial effects are due to engagement of receptor-mediated inotropic response, or due to improvement of myocardial bioenergetics.

To address these questions, we studied integrated cardiovascular response (pressure-volume analysis of the left ventricle) and isolated left ventricular (LV) trabeculae from normotensive male HanSD rats (age 20-30 weeks) after administration of 3-OHB (1 mmol/l) and drugs targeting adrenergic and cAMP-dependent signaling – metoprolol, adenylyl cyclase inhibitor 2,5-dideoxyadenosine and isoprenaline, during Tyrode solution perfusion and 1 Hz pacing.

Pressure-volume analysis demonstrated increased left ventricular end-systolic elastance (+34%), improved ventriculo-arterial coupling (+65%) and increased LV efficiency (+28%), indicating direct inotropic effect of 3OHB. In isolated LV trabeculae, 3-OHB increased twitch force in both groups, but increase was greater in healthy hearts (+55.8%) than in the HF group (+35.6%) (*p* = 0.014). The inotropic effect of 3-OHB was not further augmented by coadministration of isoprenaline and was not attenuated by beta-1 adrenergic blockade or cAMP inhibition.

Results suggest that 3-OHB directly improves LV inotropy and by a mechanism that is independent of receptor-mediated β-adrenergic signaling, most consistent with a direct bioenergetic pathway. These findings support 3-OHB as a candidate inotropic agent with a mechanistic profile distinct from conventional cAMP-dependent inotropes.

## Introduction

Ketone body 3-hydroxybutyrate (3-OHB) is produced in the liver from fatty acids (FFAs) during periods of reduced carbohydrate availability such as fasting, prolonged exercise, or caloric restriction. Under normal conditions in humans, plasma 3-OHB concentrations remain below 0.1 mM. During physiological ketosis 3-OHB levels rise to 1–2 mM, and in diabetic ketoacidosis they may exceed 10 mM.

Cardiac metabolism is dynamic and is influenced by heart failure (HF), disease severity, etiology, and comorbidities such as diabetes, obesity, or coronary artery disease. Under normal physiological conditions, FFAs are the dominant substrate, providing approximately 70-80% of energy requirements [1,2]. The failing heart exhibits reduced metabolic flexibility and impaired efficiency and increased reliance on non-FFA substrates including glucose, aminoacids and ketone bodies [3,4]. In HF, circulating levels of 3-OHB are increased due to catabolic state [5–7]. Myocardial fractional extraction of 3-OHB seems to be significantly increased in HF patients, occurring independently of systemic ketosis [8].

Interestingly, several groups observed that exogenous administration of 3-OHB in experimental animals [9] and in HF patients [10,11] increases cardiac output (CO) by up to 40%, without impairing myocardial external energy efficiency [12], in contrast to conventional inotropic agents such as dobutamine [13]. However, the extent to which this hemodynamic benefit is attributable to direct myocardial effects versus peripheral vasodilation has remained unclear. Ex vivo studies using isolated perfused Langendorff rat heart suggested direct positive inotropic effect of high dose 3-OHB (3–10 mM) [14]. The molecular mechanism underlying this direct effect remains incompletely understood. It could be due to engagement of intrinsic myocardial inotropy via to receptor mediated effects (adrenergic system or of cAMP-activation GCPR receptors), or it could be due to improvement of bioenergetics. 3-OHB may improve inotropy and energetic efficiency of the heart, potentially enhancing the amount of substrates available for oxidative cycle and contraction. The aim of this study is to characterize the direct positive inotropic effect of 3-OHB on rodent heart and to explore the mechanism of this effect by influencing adrenergic and cAMP-mediated signaling pathways.

## Methods

The study was performed and approved by the Animal Ethics Committee of IKEM and by the Ministry of Health of the Czech Republic (#12468/2021-5/OVZ). All animals used in the study were bred in IKEM, which is accredited by the Czech Association for Accreditation of Laboratory Animals.

### Pressure volume analysis

The left ventricular pressure-volume (PV) analysis was performed on normotensive male HanSD rats (n=8). Eleven-weeks male HanSD rats were anesthetized using thiopental applied intraperitoneally (80 mg/kg, VAUB Pharma a.s., Roztoky, Czech Republic). Anesthetized rat was placed on a heating pad in supine position, the ECG leads were attached to the animal (Bio Amp, ADInstruments, Dunedin, New Zeland). Tracheostomy was performed to ensure chest relaxation and to facilitate respiration during the whole operation. An intravenous cannula was inserted into the jugular vein to allow for sufficient hydration, replacement of blood loss and to administer drugs. Ballon catheter (LeMatire Single Lumen Embolectomy catheter, 2F, Burlington, MA, USA) was inserted into the left femoral vein and moved to vena cava inferior to ensure preload reductions, which is necessary for calculating load-independent measures of cardiac contractility. Pressure-volume catheter (Model SPR869, Millar, Houston, TX, USA) was placed into the LV via the right carotid artery and then connected to PowerLab (ADInstruments, Dunedin, New Zeland) and to PC with data-acquisition LabChart Pro software (ADInstruments, Dunedin, New Zeland). Echocardiography (probe 10S, 5-11.5 MHz, Vivid 7 Dimension, GE Healthcare, Chicago, IL, USA) was used to confirm correct catheter positioning throughout the procedure. Saline solution (0.5 ml) containing 3-OHB (DL-Beta-Hydroxybutyric acid sodium salt, Sigma-Aldrich, St. Louis, MO, USA) was administered intravenously. The amount of 3-OHB was calculated according to the animal’s body weight to achieve a target concentration of 1mM. After the operation, all rats were euthanized with an overdose of intravenous thiopental (200 mg/kg, VAUB Pharma a.s., Roztoky, Czech Republic) [15]. Data were recorded and analyzed by LabChart Pro software (ADInstruments, Bella Vista, NSW, Australia).

Ventricular efficiency (VE) was calculated as the ratio of stroke work (SW), defined as the area enclosed by the steady-state PV loop, to pressure–volume area (PVA): VE = SW/PVA × 100 (%). PVA was calculated as the sum of SW and potential energy, bounded by Ees, the end-diastolic pressure–volume relationship, and the systolic limb of the PV loop [16,17].

### Isolated rat ventricular trabeculae twitch force measurement

The study was performed on a group of 20-30 weeks old normotensive male HanSD rats (n=8). Animals were anesthetized with thiopental before undergoing thoracotomy and heart removal. A cardioplegic solution (5 ml Ringer’s solution 500ml with 40 ml of Thomas solution) was administered into the apex of the heart to minimize ischemia and arrest the ventricle in diastole. The heart was then removed from the pericardium and transferred onto a cold surface to minimize ischemic damage. The atria were excised, the LV was opened, and a suitable trabecula was identified and dissected. The entire heart and the trabecula from LV were weighted. The entire preparation is carried out as rapidly as possible to minimize ischemia. The trabecula was then transferred into Mayflower tissue bath (Hugo Sachs Elektronik – Harvard Apparatus GmbH, March-Hugstetten, Germany) for contractility measurement. Trabeculae were stretched to optimal length (Lmax) according to the Frank-Starling mechanism [18]. Electrical stimulation was performed by PowerLab (16/30, ADInstruments, Bella Vista, NSW, Australia) with a stimulation voltage of 10 V and duration of a single pulse 5 ms. During the measurement the perfusion solution continuously flowed through the chamber of the organ bath (6 ml/s) via a peristaltic pump (Digital Reglo, Ismatec Masterflex, Metrohm, Nederland). For the perfusion solution, we used Tyrod solution with defined concentrations (see Supplementary Table S1). It was adjusted to pH 7.40, oxygenated with carbogen (95% CO□ and 5% O□) and maintained at 37 °C using a thermostat (Optima T100, Grant Instruments, Royston, UK). The contractility force transduced via Isometric Force Transducer (FT20, Hugo Sachs Elektronik – Harvard Apparatus GmbH, March-Hugstetten, Germany) to LabChart Pro software (ADInstruments, Bella Vista, NSW, Australia).

The contractility force of trabeculae was measured at baseline at 1 Hz for 20 minutes to stabilize the twitch force (TF). Stimulation frequencies were then increased every 30 seconds in order - 0,5; 1; 2; 3; 4; 5 Hz. At each frequency the TF was measured for 30 seconds with stabilization pause of 4 minutes in between. At the end of the experiment, the length and thickness of the isolated myocardium was obtained (no trabecula was thicker than 2 mm).

We repeated the protocol after administering 1 mmol/l 3-OHB (DL-Beta-Hydroxybutyric acid sodium salt, Sigma-Aldrich, St. Louis, MO, USA) into the perfusion solution. Another measurement was obtained when isoprenaline 5µmol/l (isoprenaline hydrochloride, Sigma-Aldrich, St. Louis, MO, USA) was applied following 3-OHB administration to assess additive effects on contractile force. To determine the mechanism behind 3-OHB induced contractility we added cAMP blocker to the perfusion solution, 1 mmol/l 3-OHB and 30 µmol/l cAMP blocker (2,5 – Dideoxyadenosine, Sigma-Aldrich, St. Louis, MO, USA). Finally, we tested whether 1 mmol/l 3-OHB responds to metoprolol succinate 1 µmol/l (Sigma-Aldrich, St. Louis, MO, USA). Data were captured in LabChart Pro software (ADInstruments, Bella Vista, NSW, Australia).

We also studied group of animals (20-30 weeks old) with HF induced by permanent coronary ligation of LAD (n=8, HF) [19]. The ligation was performed at the age of 13 weeks, and echocardiography was performed to confirm reduced ejection fraction (EF) in the HF group 2 weeks after the procedure. Due to limited number of post IM animals, only responses to 3-OHB were studied.

Statistical analysis of the data was done using Graph-Pad Prism software v10.3.1 (Graph Pad Software, San Diego, CA, USA). All results are presented as a mean ± standard error of the mean. The data were analyzed by one-way ANOVA or two-way ANOVA. The values of *p* below 0.05 were considered as statistically significant.

## Results

### In vivo effect of 3-OHB on contractility of LV by pressure-volume analysis

We used a PV analysis of the LV to measure in vivo effect of 3-OHB on myocardial contractility (end-systolic elastance (Ees), preload recruitable stroke work (PRSW), coupling ratio of end-systolic elastance to arterial elastance (Ees/Ea) and ventricular efficiency (VE). The basal end systolic elastance was measured at 1.30 mmHg/µL. The administration of 3-OHB led to an increase in Ees to 1.75 mmHg/µL (change of +34%; *p* = 0.04, Figure 1b). The basal PRSW was 77.07 mmHg/µL. The administration of 3-OHB led to an increase in PRSW to 90.32 mmHg/µL (change of +17%; *p* = 0.07, Figure 1c). The basal Ees/Ea was measured at 0.35. The administration of 3-OHB led to an increase in Ees/Ea to 0.58 (change of +65%; *p* = 0.02, Figure 1d). The basal VE was 34.31%. The administration of 3-OHB led to an increase in VE to 44.03% (change of +28%, *p* = 0.02, Figure 1e).

**Figure 1.**
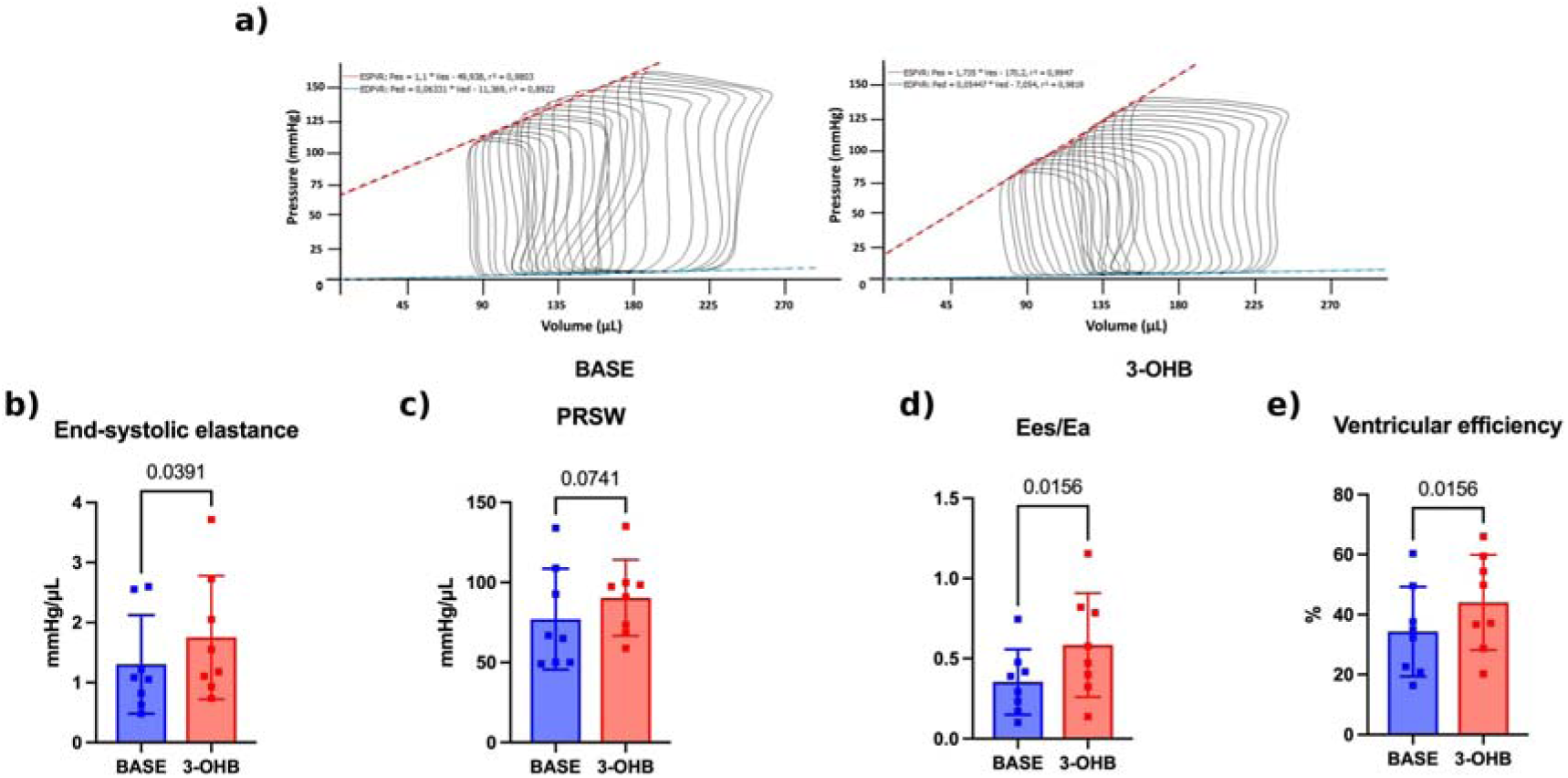
In vivo effect of 3-OHB on left ventricular contractility assessed by pressure– volume analysis. (a) Representative left ventricular pressure–volume loops recorded during inferior vena cava occlusion at baseline (BASE, left) and after intravenous 3-OHB (right); dashed lines show the end-systolic and end-diastolic pressure–volume relationships (labelled ESPVR and EDPVR in the panel); the slope of the end-systolic relationship is the end-systolic elastance (Ees). (b) End-systolic elastance (Ees). (c) Preload recruitable stroke work (PRSW). (d) Ventriculo-arterial coupling ratio (Ees/Ea). (e) Ventricular efficiency (VE). Individual values with mean ± SEM; p values are shown above the brackets.

### The impact of 3-OHB on Twitch force in healthy and in post-IM HF animals

By measuring isolated trabeculae from LV of rats, we observed a significant increase in contractile force compared with baseline conditions in healthy group. The mean contractile force rose from 0.86 cN at baseline to 1.34 cN following 3-OHB exposure (+55.8% change; *p* < 0.0001; Figure 2b). In the HF group, mean EF of infarcted rats was 24.6%, the mean TF rose from 0.59 cN at baseline to 0.80 cN after 3-OHB administration (+35.6% change; *p* = 0.0071; Figure 2b). The baseline twitch force was greater in the healthy group than in the HF group: 0.86 cN in healthy group versus 0.59 cN in HF group, the mean difference being 0.27 cN; change of 45.8%; *p* = 0.002. 3-OHB influenced TF was greater in healthy group as well: 1.34 cN versus 0.80 cN in HF group. The mean difference being 0.54 cN; change of 67.5% (*p* < 0.0001; Figure 2b). 3-OHB produced a greater improvement in TF in healthy group (+55.8%) versus in HF group (+35.6%), *p* = 0.014.

**Figure 2.**
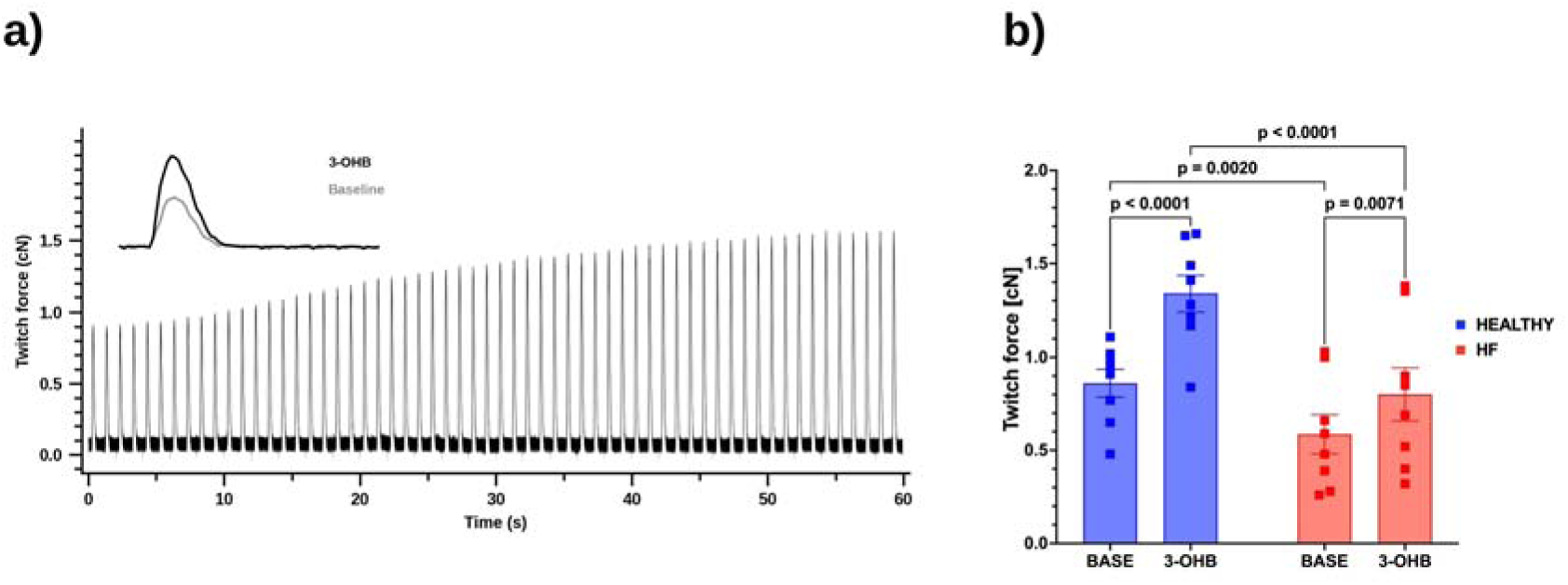
Effect of 3-OHB on twitch force of isolated left ventricular trabeculae in healthy and post-infarction heart failure rats. (a) Representative recording of twitch force during 1 Hz pacing, showing the progressive rise in developed force after 1 mmol/l 3-OHB was added to the perfusion solution. Inset: superimposed single twitches at baseline (grey) and after 3-OHB (black). (b) Twitch force at baseline (BASE) and after 3-OHB in healthy animals (blue, n = 8) and in animals with post-infarction heart failure (HF, red, n = 8). Individual values with mean ± SEM; p values are shown above the brackets.

### The effect of isoprenaline, metoprolol and cAMP inhibition on 3-OHB induced increase in TF

The baseline TF was measured at 0.39 cN in the healthy group. The administration of 3-OHB led to an increase in contractile force 0.58 cN (change of +48.7%; *p*=0.0006). Metoprolol did not produce any significant inhibitory effect, with the mean force at 0.61cN. This represents a negligible change of +5.2% compared to 3-OHB alone (*p*=0.6391, Figure 3a).

**Figure 3.**
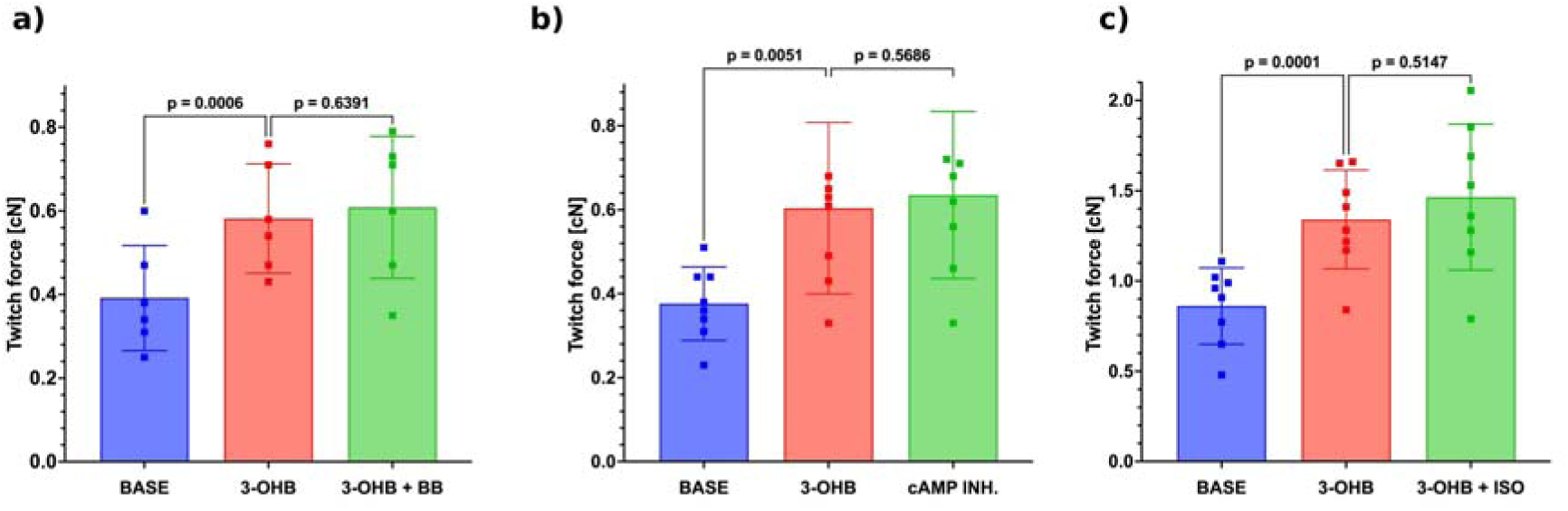
Effect of beta-1 adrenergic blockade, adenylyl cyclase inhibition and beta-adrenergic stimulation on the increase in twitch force induced by 3-OHB, in isolated left ventricular trabeculae of healthy rats. (a) Metoprolol succinate 1 µmol/l added after 1 mmol/l 3-OHB (3-OHB + BB). (b) The adenylyl cyclase inhibitor 2,5-dideoxyadenosine 30 µmol/l added after 3-OHB (cAMP INH.). (c) Isoprenaline 5 µmol/l added after 3-OHB (3-OHB + ISO). All measurements were made during 1 Hz pacing. Individual values with mean ± SEM; p values are shown above the brackets.

The mean baseline TF was 0.38 cN and after 3-OHB administration rose to 0.60 cN (a change of 57.9%, *p*=0.0051). The subsequent addition of a cAMP inhibitor resulted in a mean force of 0.63 cN, the difference compared to only 3-OHB influenced TF was + 0.03 cN (a change of +5%, *p*=0.57, Figure 3b).

The baseline twitch force was measured at 0.86 cN in the healthy group. The administration of 3-OHB led to an increase in contractile force 1.34 cN. Mean difference was 0.48 cN (a change of +55.8%; *p*=0.0001). Isoprenaline admission produced a change of 0.12 cN (+9.7%, *p=*0.51, Figure 3c).

Since we used 1 Hz as our primary studied frequency, force–frequency relationships across the full stimulation range (0.5–5 Hz) are presented in Supplementary Figures S1a-S1f.

## Discussion

The present study shows that 3-OHB has a direct positive inotropic effect on the LV myocardium, demonstrated in vivo by PV analysis and ex vivo in isolated LV trabeculae. The ex vivo effect was significant in both the healthy group and the group with HF. In healthy group occurred independently of beta-adrenergic and cAMP-mediated stimulation, supporting direct effect of 3-OHB on bioenergetics.

The PV analysis reflects the effect of 3-OHB on the whole cardiovascular system. Here the dominant change was an increase in contractility rather than a reduction of afterload. Ees a load-independent measure of intrinsic contractility, rose by 34% (*p* = 0.04), with a parallel trend in preload recruitable stroke work (+17%, *p* = 0.07). Ventriculo-arterial coupling (Ees/Ea) improved by 65% (*p* = 0.02). Because Ees/Ea can improve either through a higher Ees or through a lower arterial elastance, the fact that Ees itself increased places the improvement mainly in the ventricle rather than in the periphery. VE rose by 28% (*p* = 0.02), meaning that more external work was produced for the same energy expenditure, indicating improved mechanic efficiency.

In isolated trabeculae 3-OHB increased twitch force by 55.8% (*p* < 0.0001) in healthy myocardium. The similar effect was present after MI. In trabeculae from rats with established MI (mean EF 24.6%), 3-OHB increased twitch force by 35.6% (*p* = 0.0071). Absolute force was lower than in healthy controls, but the diseased, remodeled myocardium still responded to 3-OHB.

Interestingly, we observed enhancement of inotropy not only in disease, but also even in normal non-diseased myocardium, indicating presence of contractile reserve recruitable by 3-OHB. Because 3-OHB levels and flux increase in rodents during acute exercise [20], the 3-OHB-induced increase in contractility may be a part of the physiological response of the heart to increased stress, acting in parallel with, but independently of, sympathetic stimulation.

To at least partly differentiate between bioenergetic and receptor-mediated pathways, we used co-administration of metoprolol with 3-OHB to assess the potential involvement of beta-adrenergic receptor activation. The absence of any significant change in contractile force following addition of metoprolol to the solution with 3-OHB (change of +5.2%, *p*=0.6391) indicates that the inotropic effect of 3-OHB is independent of beta-1 adrenergic receptor stimulation.

Furthermore, we used 2,5-dideoxyadenosine (DDA), an inhibitor of adenylyl cyclase, which blocks the enzyme non-competitively with an IC of approximately 3 µM (in our protocol we used 30 µM) [21]. It is important to note that DDA primarily attenuates beta-agonist-stimulated cAMP accumulation rather than abolishing basal intracellular cAMP levels. Its inhibitory potency is therefore concentration-dependent and partial, we used concentration which blocks more than 90% of receptors. Co-administration of DDA with 3-OHB produced no significant change in twitch force compared to 3-OHB alone (baseline of 0.38, after 3-OHB increase to 0.60 cN a change of 57.9%, *p*=0.0051), suggesting that the inotropic effect of 3-OHB does not depend on receptor-stimulated cAMP signaling. While a residual contribution of basal cAMP cannot be entirely excluded with this experimental approach, these data argue against a major role of Gαi-coupled receptor activation in the observed inotropy.

To discover potential additional effect of B1AR stimulation on top of 3-OHB, we used isoprenaline TF increased only slightly and non-significantly (+9.7%, *p* = 0.51), indicating that inotropic response after 3-OHB may be submaximal and cannot by further enhanced by additional B1AR stimulation. Because the study was not powered to detect an effect of this size, we cannot determine whether 3-OHB and beta-1 adrenergic stimulation act through separate, additive pathways or whether the response to 3-OHB already approaches the maximal contractile capacity of the preparation.

Probable explanation for the observed positive inotropy, given the absence of receptor-mediated cAMP signaling and adrenergic stimulation, is a direct bioenergetic effect. Under conditions of stable extracellular calcium concentration (as maintained in our Tyrode perfusion solution), additional ATP availability could support enhanced cross-bridge cycling and accelerated SR Ca² re-uptake via SERCA, thereby increasing twitch amplitude without a primary change in calcium transient amplitude — consistent with the observation in cardiac organoids that 3-OHB increased contraction amplitude without affecting calcium handling or conduction velocity [22]. The presence of 10 mM glucose in the perfusion solution throughout the experiment is an important consideration: since contractile force increased despite adequate glucose supply, the inotropic effect of 3-OHB is unlikely to reflect a substrate substitution in a substrate-depleted preparation. Rather, it suggests a true additive bioenergetic contribution of 3-OHB on top of existing glucose oxidation.

### Clinical perspective

From a clinical perspective, the results of this study provide mechanistic support for 3-OHB as a candidate inotropic agent. Currently available positive inotropes (catecholamines and PDE3 inhibitors), improve hemodynamics but are associated with increased mortality in chronic use, partly due to their dependence on cAMP-mediated pathways, which accelerate oxygen consumption, worsen efficiency in an already energy-depleted failing heart [23] and myocardial remodeling and fibrosis by activating profibrotic pathways [24,25]. The mechanism of 3-OHB, as suggested by this study, appears to be different: it increases contractile force through a metabolic pathway [26]. This metabolic safety profile distinguishes 3-OHB from existing inotropes and warrants further clinical investigation.

### Limitations

The absence of cross-sectional area normalization of trabeculae force represents a methodological limitation: absolute twitch forces between animals and groups are not directly comparable without accounting for differences in muscle geometry. Passive tension was not systematically measured, which may have introduced variability in the assessment of optimal sarcomere length and the Frank-Starling relationship. Due to limited number of post-IM animals, we did not study pharmacological interventions on top of 3-OHB in this group. The respiratory state of isolated trabeculae cannot be directly monitored in this preparation, and gradual hypoxia cannot be excluded over long experimental protocols. Variability in perfusate pH may have introduced minor beat-to-beat variability in contractile measurements. Finally, the surgical technique for heart explantation introduces an unavoidable ischemic interval, which may affect trabecular viability. However, the use of cardioplegic arrest and cold transfer minimizes this effect.

## Conflict of interest

None declared.

## Grant support

Supported by Ministry of Health, Czech Republic - conceptual development of research organization („Institute for Clinical and Experimental Medicine – IKEM, IN 00023001“) and by the Ministry of Health of the Czech Republic in cooperation with the Czech Health Research Council within the National Institute CarDia, project No. NW26A-CARDIA.

## Declaration on the Use of Artificial Intelligence

The authors used Anthropic’s Claude Opus 5 to edit Figure 2a. All outputs were reviewed and verified by the authors for factual accuracy.

---

**Supplementary Figure S1.** Effect of 3-OHB on twitch force of isolated left ventricular trabeculae across the stimulation frequency range. Twitch force at baseline (BASE) and after 1 mmol/l 3-OHB in healthy animals (blue) and in animals with post-infarction heart failure (HF, red), at (a) 0.5 Hz, (b) 1 Hz, (c) 2 Hz, (d) 3 Hz, (e) 4 Hz and (f) 5 Hz. Individual values with mean ± SEM; p values are shown above the brackets.

**Supplementary Table S1.** Composition of the Tyrode perfusion solution used for the isolated left ventricular trabeculae experiments.

## Notes

### Competing Interest Statement

The authors have declared no competing interest.

## References

1. Bing RJ, Siegel A, Ungar I, Gilbert M. Metabolism of the human heart: II. Studies on fat, ketone and amino acid metabolism. Am J Med 1954;16:504–515.

2. Murashige D, Jang C, Neinast M, Edwards JJ, Cowan A, Hyman MC, Rabinowitz JD, et al. Comprehensive quantification of fuel use by the failing and nonfailing human heart. Science 2020;370:364–368.

3. Lopaschuk GD, Karwi QG, Tian R, Wende AR, Abel ED. Cardiac energy metabolism in heart failure. Circ Res 2021;128:1487–1513.

4. Neubauer S. The failing heart – an engine out of fuel. N Engl J Med 2007;356:1140–1151.

5. Lommi J, Kupari M, Koskinen P, Näveri H, Leinonen H, Pulkki K, Härkönen M. Blood ketone bodies in congestive heart failure. J Am Coll Cardiol 1996;28:665–672.

6. Melenovsky V, Benes J, Franekova J, Kovar J, Borlaug BA, Segetova M, Tura A, et al. Glucose homeostasis, pancreatic endocrine function, and outcomes in advanced heart failure. J Am Heart Assoc 2017;6:e005290.

7. Monzo L, Kovar J, Borlaug BA, Benes J, Kotrc M, Kroupova K, Jabor A, et al. Circulating beta-hydroxybutyrate levels in advanced heart failure with reduced ejection fraction: determinants and prognostic impact. Eur J Heart Fail 2024;26:1931–1940.

8. Monzo L, Sedlacek K, Hromanikova K, Tomanova L, Borlaug BA, Jabor A, Kautzner J, et al. Myocardial ketone body utilization in patients with heart failure: the impact of oral ketone ester. Metabolism 2021;115:154452.

9. Gopalasingam N, Moeslund N, Christensen KH, Berg-Hansen K, Seefeldt J, Homilius C, Nielsen EN, et al. Enantiomer-specific cardiovascular effects of the ketone body 3-hydroxybutyrate. J Am Heart Assoc 2024;13:e033628.

10. Sramko M, Wohlfahrt P, Kissimon F, Svirlochova V, Franekova J, Kautzner J, Melenovsky V. Hemodynamic effects of oral ketone ester in acutely decompensated heart failure with low cardiac output syndrome. JACC Heart Fail 2026 (in press).

11. Berg-Hansen K, Gopalasingam N, Christensen KH, Ladefoged B, Andersen MJ, Poulsen SH, Borlaug BA, et al. Cardiovascular effects of oral ketone ester treatment in patients with heart failure with reduced ejection fraction: a randomized, controlled, double-blind trial. Circulation 2024;149:1474–1489.

12. Nielsen R, Møller N, Gormsen LC, Tolbod LP, Hansson NH, Sorensen J, Harms HJ, et al. Cardiovascular effects of treatment with the ketone body 3-hydroxybutyrate in chronic heart failure patients. Circulation 2019;139:2129–2141.

13. Francis GS, Bartos JA, Adatya S. Inotropes. J Am Coll Cardiol 2014;63:2069–2078.

14. Homilius C, Seefeldt JM, Axelsen JS, Pedersen TM, Sørensen TM, Nielsen R, Wiggers H, et al. Ketone body 3-hydroxybutyrate elevates cardiac output through peripheral vasorelaxation and enhanced cardiac contractility. Basic Res Cardiol 2023;118:37.

15. Miklovic M, Kala P, Melenovsky V. Simultaneous biventricular pressure-volume analysis in rats. J Physiol Pharmacol 2023;74:131–147.

16. Suga H, Sagawa K. Instantaneous pressure-volume relationships and their ratio in the excised, supported canine left ventricle. Circ Res 1974;35:117–126.

17. Suga H. Total mechanical energy of a ventricle model and cardiac oxygen consumption. Am J Physiol 1979;236:H498–H505.

18. Dhein S. Isolated papillary muscles. In: Practical Methods in Cardiovascular Research, Dhein S, Mohr FW, Delmar M (eds), Springer-Verlag, Berlin Heidelberg, 2005, pp 190–197.

19. Selye H, Bajusz E, Grasso S, Mendell P. Simple techniques for the surgical occlusion of coronary vessels in the rat. Angiology 1960;11:398–407.

20. Axsom J, TeSlaa T, Lee WD, Chu Q, Cowan A, Bornstein M, Neinast M, et al. Quantification of nutrient fluxes during acute exercise in mice. Cell Metab 2024;36:2560–2579.

21. Hartmann M, Schrader J. Isoproterenol antagonistic effect of 2′,5′-dideoxyadenosine in the isolated perfused guinea-pig heart. J Mol Cell Cardiol 1993;25:331–338.

22. Seefeldt JM, Libai Y, Berg K, Jespersen NR, Lassen TR, Dalsgaard FF, Ryhammer P, et al. Effects of ketone body 3-hydroxybutyrate on cardiac and mitochondrial function during donation after circulatory death heart transplantation. Sci Rep 2024;14:757.

23. Belletti A, Castro ML, Silvetti S, Greco T, Biondi-Zoccai G, Pasin L, Zangrillo A, et al. The effect of inotropes and vasopressors on mortality: a meta-analysis of randomized clinical trials. Br J Anaesth 2015;115:656–675.

24. Teerlink JR, Pfeffer JM, Pfeffer MA. Progressive ventricular remodeling in response to diffuse isoproterenol-induced myocardial necrosis in rats. Circ Res 1994;75:105–113.

25. Akiyama-Uchida Y, Ashizawa N, Ohtsuru A, Seto S, Tsukazaki T, Kikuchi H, Yamashita S, et al. Norepinephrine enhances fibrosis mediated by TGF-β in cardiac fibroblasts. Hypertension 2002;40:148–154.

26. Psotka M, Gottlieb S, Francis G, Allen L, Teerlink J, Adams K, Rosano GM, et al. Cardiac calcitropes, myotropes, and mitotropes: JACC review topic of the week. J Am Coll Cardiol 2019;73:2345–2353.

