## Supplemental Table 1 for "Inotropic effects of 3-hydroxybutyrate in a rodent heart: pressure-volume and isolated rat ventricular trabeculae analysis"

**Supplementary Table S1.** Composition of the Tyrode perfusion solution used for the isolated left ventricular trabeculae experiments.

| **Perfusion solution** | **Concentration (mmol/l)** |
| --- | --- |
| NaCl | 137 |
| D-glucose | 10 |
| HEPES | 5 |
| KCl | 4.5 |
| MgCl₂ | 1 |
| NaOH | 2 |
| CaCl₂ | 2 |

The solution was adjusted to pH 7.40, oxygenated with carbogen and maintained at 37 °C.
