## Supplementary figures and images for "Inotropic effects of 3-hydroxybutyrate in a rodent heart: pressure-volume and isolated rat ventricular trabeculae analysis"

### Supplementary Figure 1

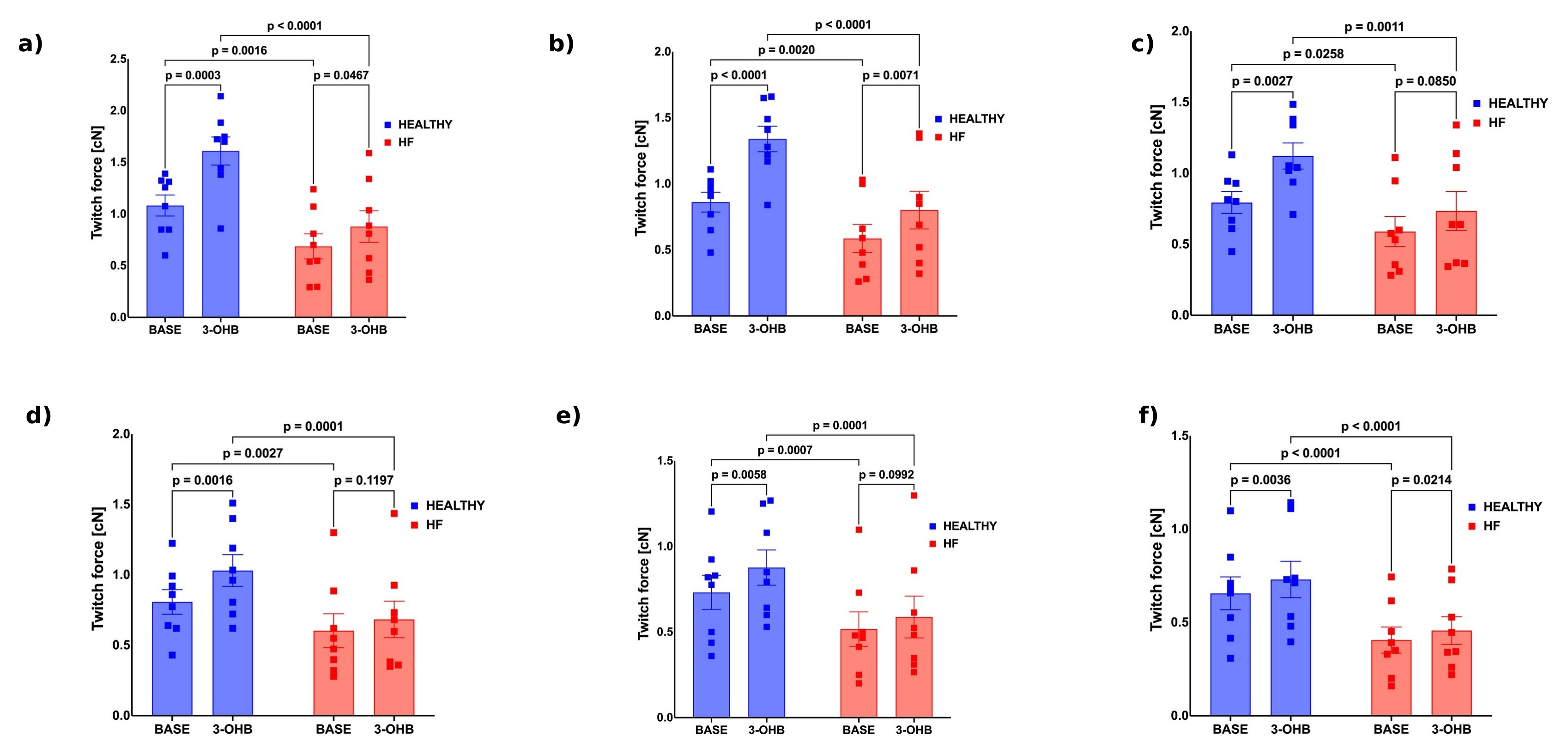
